# Modelling Audio-Visual Reaction Time with Recurrent Mean-Field Networks

**DOI:** 10.1101/2025.05.29.656149

**Authors:** Rebecca M. Brady, John S. Butler

## Abstract

Audio-visual stimuli are typically detected faster and more accurately than unisensory ones, but it remains unclear whether this benefit reflects true integration of evidence or other effects such as independent processing and modality repetition. We address this with three biologically motivated models of audio-visual integration, built on Wong and Wang model, a mean-field recurrent framework originally developed for unisensory decision-making.

The OR model takes in independent unisensory processes through winner-take-all competition, and the SUM model implements full linear summation of sensory evidence. These two sit at opposite ends of the integration spectrum. The REPEAT model adds switch and repeat costs, whereby responses are faster when the modality repeats across trials. We simulated 121 participants whose unisensory evidence accumulation rates varied from modality dominance to balanced integration. We fitted the simulated reaction times and accuracy with a drift diffusion model and compared the resulting multisensory drift rates with the optimal prediction from the unisensory rates, as well as with empirical audio-visual detection data. The OR and SUM Models reproduced established unisensory response patterns from behavioural data from in children, children with autism, adults, older adults, and people with Parkinson’s disease. Drift diffusion analysis revealed suboptimal integration in the OR Model and optimal integration in the SUM Model. However, the SUM Model also produced supra-optimal responses under certain conditions, inconsistent with behavioural data. The REPEAT Model captured the role of priming in sensory repetition effects, distinguishing it from true multisensory integration.

Together, these models highlight how biologically grounded mathematical frameworks can shed light on the mechanisms underlying multisensory integration, particularly the nuanced contributions of modality repetition and integration efficiency.

## Introduction

Multisensory integration is essential for how we perceive and interact with our environment, enhancing detection and reaction time to stimuli and improving cognitive and behavioural performance (1–5). The ability to integrate signals develops during childhood, progressing from no integration to the combination of signals (6–8), and this trajectory is often delayed in individuals with autism (9, 10). In contrast, older adults show atypical multisensory behaviour, such as an increased susceptibility to perceptual illusions, which predicts falls (11–14). Despite its importance, the mechanisms underlying these behavioural differences remain unclear, and theoretical models can help to address them (15, 16).

A central question is whether the speed-up in responses to multisensory stimuli reflects true integration of sensory evidence or can be explained without it. Independent-channel (race) models predict a benefit from selecting the faster of two unisensory responses on each trial, and violations of the race-model bound have been taken as evidence of integration (17). Alternatively, integration can be framed in terms of optimal cue combination, in which the multisensory drift rate is predicted from the unisensory rates (18). Recent work suggests that part of the observed benefit may arise from factors other than integration, such as switch and repeat effects across consecutive modalities (16, 19). This leaves open the question of how these accounts can be compared within a single, biologically grounded framework that simulates trialby-trial behaviour at both individual and group levels.

Here we address this using the mean-field recurrent model of (20), which describes the population dynamics of area LIP during perceptual decision-making and has been used to link electrophysiological and behavioural findings (21). The model includes noise to produce trial-to-trial variability, and its parameters allow sensitivity to vary across individuals, which we use to simulate auditory or visual dominance. We extend this unisensory model to three models of audio-visual integration and simulate a seminal audio-visual detection task using congruent, salient stimuli, which has been used with children (6), children with autism (9), adults (1, 3), older adults, and people with Parkinson’s disease (22).

The ***OR Model*** implements a non-integrative Winner-Take-All (WTA) strategy in which the faster of the auditory and visual unisensory processes determines the multisensory response. It is motivated by developmental data showing that segregated processing is consistent with behaviour in younger children and children with autism (6, 9, 23), and it provides the baseline: behaviour that exceeds its predictions cannot be explained by independent processing alone. The ***SUM Model*** implements linear summation of evidence, with distinct auditory, visual and integrative areas. This is motivated by the multisensory benefits observed in adults (1, 3, 13, 22, 24) and by evidence that additive neural activity in mouse cortex matches additive behavioural measures (25). The ***REPEAT Model*** extends the OR Model with priming when the modality repeats across trials, following (19, 23), to test whether segregated processing with sequence effects can account for the apparent benefit.

To compare the models with behaviour, we simulate participants with varying unisensory sensitivities and compare the output with published data from neurotypical children and children with autism (9), people with Parkinson’s disease and age-matched controls (22), and adults (19). We fit reaction times and accuracy jointly with a drift-diffusion model (26), and compare the multisensory drift rates with the optimal prediction from the unisensory fits (18). We hypothesised that the OR Model would be suboptimal, that the SUM Model would approach optimal integration, and that the REPEAT Model would reproduce repetition effects without integration.

## Methods

### Mathematical Model

The reduced two-variable model is a mean-field model designed by (20) which is a system of non-linear ordinary differential equations that describe the dynamics of the accumulation of competing activity between two populations of cortical neurons for unisensory decision-making. Our multisensory adaptation of this framework consists of populations of neurons divided into distinct sensory columns, representing auditory (blue) and visual (red) and audio-visual (gray) areas (Figure 2).

**Fig 1.**
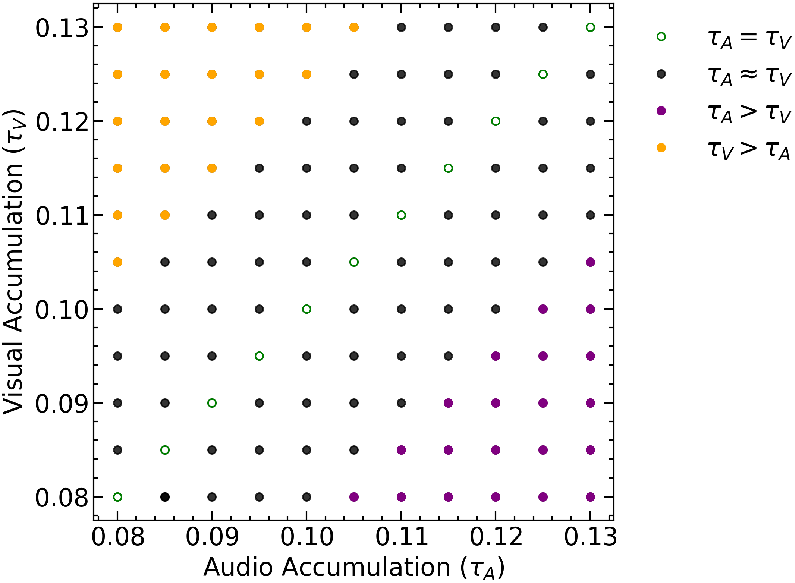
All possible combinations of values for the audio and visual accumulation, *τ*_*A*_ and *τ*_*V*_ . Each dot represents one simulated participant and simulates variability in audio and visual sensitivities across human participants. The green dots along the diagonal represent simulated participants with equal audio and visual sensitivity values. While the purple and orange dots are for large disparities between the unisensory accumulation. A grey version of the grid is superimposed on the contour plots to highlight the effect of unisensory accumulation on multisensory outcomes.

**Fig 2.**
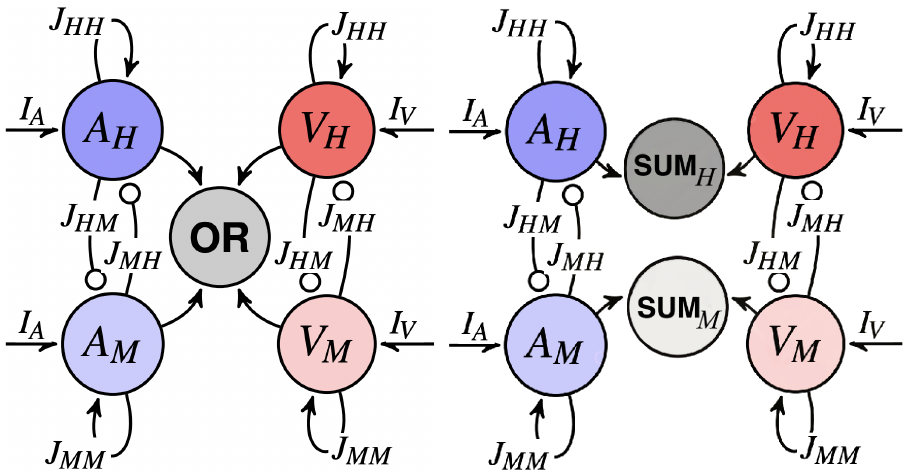
Left: The OR Model architecture with distinct audio (A) and visual (V) ‘Hit’ (H) and ‘Miss’ (M) populations. Arrows show excitatory connections, while lines with open circles show inhibitory connections. When activity in any of the neural populations (*A*_*H*_, *A*_*M*_, *V*_*H*_, or *V*_*M*_ ) reaches a threshold value, the reaction time and accuracy are recorded in the ‘OR’ variable. Right: The linear summation SUM Model architecture with distinct unisensory audio (A) and visual (V) areas and a multisensory summation (SUM) area. Each sensory region comprises two ‘Hit’ (H) and ‘Miss’ (M) populations. Arrows show excitatory connections, while lines with open circles show inhibitory connections. The audio (blue) and visual (red) components receive sensory input (*I*_*V*_ and *I*_*A*_) and combine their accumulations in the multisensory area (grey) as a linear summation. The reaction time and accuracy are recorded when activity in either the *SUM*_*H*_ or *SUM*_*M*_ neural populations reaches a threshold value.

Each neural population in the models is self-excitatory and extra-inhibitory, and all build their firing rate in response to an external input as time passes in the system.

The population activity in the system can be described entirely by the following Equations (1) - (6), simulated for four neural units (*A*_*H*_, *A*_*M*_, *V*_*H*_, or *V*_*M*_ ) in two distinct audio and visual areas (*S* ∈ *{A, V }*), each with two competing ‘Hit’ and ‘Miss’ neural populations (*i* ∈ *{H, M }*) in the detection task,

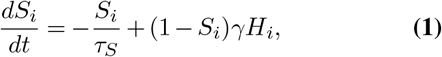

where the kinetic parameter *γ* = 0.641 and *τ*_*S*_ is a time constant associated with modality *S* which governs the sensitivity and accumulation rate of information, this parameter will be systematically varied to simulate different individual participants reaction times and accuracy from a range of populations from development to old age.

The population firing rate, *H*_*i*_, is a function of the input and output currents defined by *x*_*H*_ and *x*_*M*_ . The parameters *a* = 270 V nC*−*1, *b* = 108 Hz, and *d* = 0.154 s correspond to the gain factor, threshold potential, and noise factor, respectively,

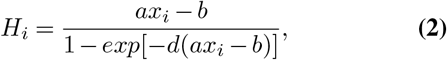

where,

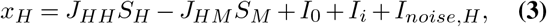

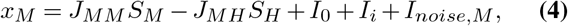

*x*_*H*_ and *x*_*M*_ describe the total synaptic current received by each population as a sum of all recurrent inputs and inputs external to the system. Recurrent inputs accrue from activity in connective populations. They are governed by the strength of the connection from population *i* to population *j* is described by *J*_*i,j*_ for *i, j* ∈ {*H, M*} where populations are self-excitatory and other inhibitory, *J*_*HH*_ = *J*_*MM*_ = 0.2609 nA and *J*_*HM*_ = *J*_*MH*_ = 0.0497 nA.

There are three input variables external to the system. *I*_0_ = 0.3255 nA is a common external input to both populations. *I*_*i*_, described by Equation 5, encodes stimulus information and is received by the ‘Hit’ and ‘Miss’ populations within corresponding modalities. It represents information received by the LIP populations from earlier stimulus encoding areas such as the middle temporal area for visual encoding and the temporoparietal cortex or superior colliculus for auditory encoding (27). The strength of these upstream connections is modulated by *J*_ext_ = 5.2 × 10*−*5 nA Hz*−*1. The strength of *I*_*i*_ depends on the stimulus strength *c*, which has been set as *c* = 12% to ensure a stable system with high accuracy just below the ceiling level, producing outputs consistent with the audio-visual literature with a mean firing rate of *µ*_0_ = 30 Hz,

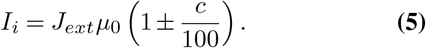

Time-varying noise, *I*_*noise,i*_ (Equation 6), is added as a Brownian motion process (28) with Gaussian white noise, *η*, and standard deviation *σ*_noise_ = 0.007 nA modulated by the AMPA time constant *τ*_AMPA_ = 2 ms,

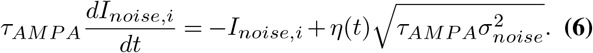

For each simulation when the firing rate reaches a threshold of 20 Hz the reaction time and response (Hit or Miss) are recorded. The inclusion of time-varying noise enables the stimulation of trial by trial reaction time and response variability.

The models code were written in Python (29). The models were numerically integrated with the 4th Order Runge-Kutta method with a time-step of 0.1 ms in Python using Numpy and Matplotlib packages (30–32).

### Individual Simulated Participants

Each of the models had 121 simulated participants with distinct sensitivity to audio and visual stimuli, which were modelled by unique combinations of sensory accumulation values for *τ*_*A*_ and *τ*_*V*_ :

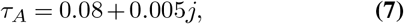

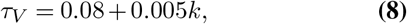

for *j, k* ∈ (0, 1, 2, 3, 4, 5, 6, 7, 8, 9, 10), resulting in combinations of *τ* values (Figure 1). Higher *τ* values denote faster accumulation of sensory information in the corresponding sensory modality (purple and orange dots). The selection of *τ* values aims to ensure the biological plausibility of reaction time outcomes in the unisensory responses. Figure 1, shows the values of audio (*τ*_*A*_) and visual (*τ*_*V*_ ) accumulations used. The green data points along the diagonal signify simulated participants with equivalent sensitivities to both modalities, thus exhibiting comparable reaction times on both auditory and visual trials.

The grey off-diagonal data points are simulated participants with slightly faster accumulation rates in one modality relative to the other. Specifically, simulated participants positioned the upper diagonal exhibit more rapid responses to visual stimuli (orange), whereas lower diagonal display heightened sensitivity to auditory stimuli (purple). Instances where simulated participants deviate markedly from the diagonal reflect significant disparities in processing speed between modalities. Disparities in modality sensitivity are commonly observed in biological participants, although wider discrepancies are potentially indicative of deficiencies in the processing of the slower modality. Within this framework, simulated participants with notably low *τ* values in one modality essentially function as unisensory responders.

For the OR and SUM Models each simulated participant had 3000 trials equally divided across the modalities (1000 audio, 1000 visual, and 1000 audio-visual). For the REPEAT model each simulated participant had 4500 trials, comprising of 1500 trials per sensory modality with 750 switch and 750 repeat trials.

### Multisensory Models

Figure 2 shows the architecture of the OR Model (left) and SUM Model (right). The auditory (blue) and visual (red) populations, each receiving sensory input (*I*_*V*_ and *I*_*A*_) and featuring self-excitatory connections (*J*_*HH*_ and *J*_*MM*_ ) and within modality inhibition (*J*_*HM*_ and *J*_*MH*_ ) between the ‘Hit’ (H) and ‘Miss’ (M) neural populations.

### OR Model

In the OR Model the individual trial outcomes are exclusively attributed to either an auditory or visual population. Upon the activation levels of the first of the neural populations (*A*_*H*_, *A*_*M*_, *V*_*H*_, or *V*_*M*_ ) reaching a predetermined threshold value of 20 Hz, as depicted in Figure 3A, the choice (Hit or Miss) and reaction time are recorded. Any behavioural benefit observed in outcomes derived from the Winner Take All (WTA) approach used in this OR Model stems from averaging the proportion of correct choices or reaction times across multiple trials, Figure 2.

**Fig 3.**
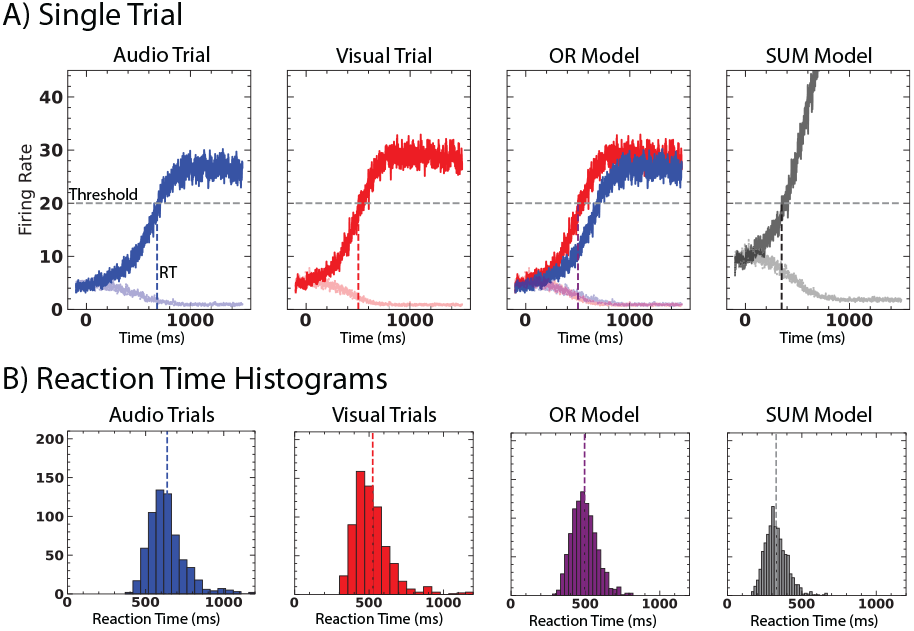
Simulated Participant Data (A) Simulations of neural population activity dynamics for one simulated participant on a single trial, which includes an audio-alone condition (left), visual-alone condition (middle left), the audio-visual OR Model condition (middle right) and the audio-visual SUM Model condition (right). Dark blue accumulations show the audio hit population, while light blue shows the audio miss population. Dark red accumulations show the visual hit population, while light red shows the visual miss. Dark grey accumulations show the SUM Model hit accumulations, and light grey show the SUM Model miss accumulations. The horizontal dashed line indicates the threshold activity level. The vertical dashed line shows the time the threshold is first reached, and reaction time and accuracy are recorded. (B) Reaction time histograms for one simulated participant across all trials. This simulated participant had similar reaction time distributions for the audio- and visual-alone conditions. The vertical dashed lines indicate the mean value of the distribution.

### SUM Model

The Linear Summation model (SUM Model) is informed by a common multisensory integration strategy, which posits that sensory information from multiple modalities is combined and processed with each sensory modality, contributing to a unified perceptual experience Figure 3B. The SUM Model reaction times as a co-activation account of multisensory processing which has been observed to accurately depict reaction time data across many audio-visual studies (1, 25, 33, 34).

The outputs of these sensory populations feed into a decision-making area where ‘Hit’ and ‘Miss’ accumulations from the audio and visual areas combine in their respective ‘SUM’ units (Figure 2 (grey)). Within these units, the reaction time and choice (Hit or Miss) for each condition (A, V, and AV) are recorded when one unit reaches a threshold value (Figure 3). The unisensory outputs are combined in a linear summation. In the SUM Model, sensory information from auditory and visual modalities is independently gathered in two separate sensory areas, each featuring recurrent connections specific to its modality. The crux of the SUM Model lies in a decision-making area where audio and visual inputs converge, and a unified perceptual experience is formed. Unlike the previously presented OR Model, the decision-making area comprises two distinct, unconnected populations: one responsible for processing Hit accumulations (*SUM*_*H*_ ) and another for processing Miss accumulations (*SUM*_*M*_ ). A decision is reached when the Hit or Miss population reaches a threshold value of 20 Hz. Figure 2 describes the interconnected structure of the SUM Model, illustrating the convergence of modality-specific information through a linear summation process.

### REPEAT Model

The REPEAT Model represents an extended version of the OR Model. The REPEAT model accounts for sensory priming and task-switching effects by first categorising trials as either ‘switch’ or ‘repeat’. The switch and repeat category assigned to the current trial depends on the previous trial (19). Repeat trials occur when the sensory condition in the current trial has also been presented in the previous trial;

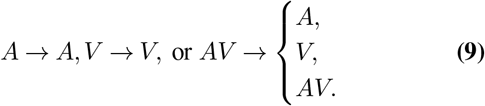

Switch trials occur when a condition in the current trial is absent from the condition presented in the previous trial;

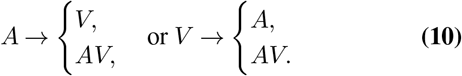

The REPEAT Model implements an increase to the system’s initial conditions during repeat trials to replicate the effects of sensory priming. During these repeat trials, the model exhibits a baseline activity increase of one standard deviation of the noise to the ‘Hit’ neural population (*A*_*H*_, *V*_*H*_ ) of the repeated (Equation 1). In *AV* → *AV* trials, the audio and visual hit populations receive an activity increase of one standard deviation of the noise, (35, 36).

#### Behavioural Analysis

The three models (OR Model, SUM Model, and REPEAT model) generate reaction time and accuracy data for audio and visual unisensory and multisensory audio-visual trials. Each condition (A, V, and AV) consists of a ‘Hit’ or ‘Miss’ neural unit, which accumulates activity and relates to detecting or missing sensory input. For each condition, reaction time and accuracy are recorded based on which of the two neural units is first to reach a threshold value of 20 Hz (Figure 3). Individual simulated participants mean reaction times and accuracy were submitted to a one-way analysis of variances (ANOVAs), with the factor of stimulus modality (audio, visual and audio-visual) with a factor of model-type (OR or SUM) for the model comparison and a factor of trial-type (Repeat or Switch) for the REPEAT model, with follow-up post hoc Tukey’s Honestly Significant Difference (HSD) test.

### Multisensory Analysis

To investigate the multisensory responses, the individual simulated participant reaction time and accuracy data (26) for the three sensory conditions (A, V, and AV) were fit to a general drift-diffusion model (GDDM) (37). The GDDM consisted of three bounded free parameters, non-decision time (*t*_*nd*_ ∈ [0, 800] ms), noise (*σ* ∈ [0.5, 2.5]), and drift rate, (*r*_*i*_ ∈ [1, 25]), for the audio, visual, and audio-visual conditions. Mean simulated participant drift rates and non-decision times were submitted to a one-way analysis of variances (ANOVAs), with the factor of stimulus modality (audio, visual and audio-visual) for all models and a factor of trial type (Repeat or Switch) for the REPEAT model, with follow up post-hoc paired t-tests for all. The observed audio, *r*_*A*_, and visual, *r*_*V*_, drift rates were used to calculate the optimal audio-visual drift rate predicted,

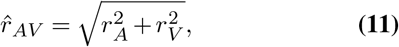

which is our metric for evaluating optimal multisensory integration (18). The observed multisensory drift rate *r*_*AV*_ and the predicted drift rate 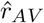 from Equation 11 were submitted to a paired t-test and a correlation analysis.

## Results

### OR Model and SUM Model Results

#### Single Simulated Participant’s Behavioural Results

Both the OR Model and SUM Model were applied to 121 unique simulated participants, each with different paired combinations of sensitivity to audio and visual stimuli, with 3000 trials over the Audio, Visual, and Audio-Visual conditions. Figure 3 presents example trials from the auditory (left), visual (middle left), audio-visual OR Model (middle right), and audio-visual SUM Model (right), depicting how neural activity accumulates across populations in each case. The ‘Hit’ population activities are the darker colours, while the ‘Miss’ population activities are the lighter colours. In the trial shown in Figure 3, activity in the ‘Hit’ populations on all trials reached the threshold value, indicating that a stimulus was detected (i.e. a ‘Hit’ result). Reaction time and accuracy are determined by the first accumulation to reach the threshold in the trials illustrated in Figure 3, that is, the activity of the auditory ‘Hit’ population. Figure 3 (bottom) shows the distribution of reaction times for a simulated participant across the audio, visual and audio-visual trials. As expected, the mean value and the spread of the audio-visual reaction times are faster than those of the unisensory reaction times and the mean of the SUM Model trials is faster than that of the OR Model.

#### Group Behavioural Results

The OR Model (left column) and SUM Model (right column) simulated participants’ reaction times (row) and accuracy (bottom row) are depicted in Figure 4. The plots show the audio-alone condition (blue), visual-alone condition (red), and audio-visual condition (grey), with each point representing a simulated participant’s average reaction time and accuracy.

**Fig 4.**
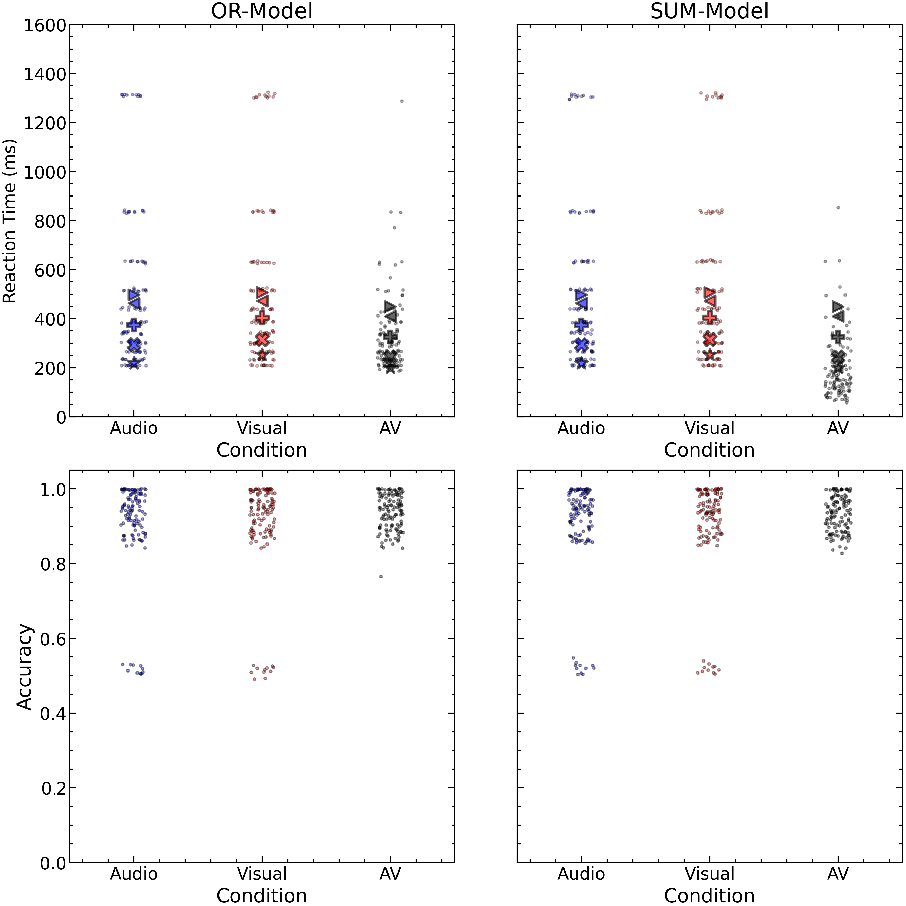
Reaction times (top) and Accuracy (bottom) per sensory condition for the OR Model (left) and SUM Model (Right). Each dot represents a simulated participant. Other symbols show average reaction time data on audio-visual detection tasks with the following participants: children with autism spectrum disorder (►) and age-matched controls (◄) from (9), older adults with Parkinson’s Disease (+) and age-matched healthy controls (X) from (22), and healthy adults (*) from (19).

The stratification seen in unisensory reaction time plots reflects the distribution of *τ* values that define audio and visual sensitivities across simulated participants. As shown in Figure 1, 121 unique *τ*_*A*_ and *τ*_*V*_ combinations were tested, with 11 simulated participants exhibiting similar unisensory behaviour (with minor variation). This stratification is less apparent in the audio-visual reaction times. Audio-visual trials show faster reaction times and accuracy for both the OR and SUM Models. To compare simulated reaction times with experimental data, Figure 4 (left) shows group average reaction times from audio-visual detection studies. From top to bottom, these symbols represent the average reaction times for children with autism spectrum disorder (►), their age-matched controls (◄) (9), older adults with Parkinson’s Disease (+), their age-matched healthy controls (X) (22), and healthy adults (*) (19). The simulated participants unisensory data spans a similar range to the experimental unisensory data. The OR Models audio-visual reaction times overlap with the experimental data while for the SUM Model there are a number of the simulated participants whose reaction times were faster than those of observed in experimental studies.

For the reaction time data a 2 (Model: OR, SUM) × 3 (Condition: Audio, Visual, Audio-Visual) repeated-measures ANOVA was conducted. The analysis revealed significant main effects of Model, F(1, 120) = 4.615, p< 0.05, and Condition, F(2, 240) = 59.87, p< 0.01, as well as a significant Model × Condition interaction, F(2, 240) = 4.62, p< 0.05. Tukey HSD post hoc tests showed Audio-Visual conditions resulted in significantly lower reaction time than both the Audio (M difference = 232.5, p< 0.01) and Visual conditions (M difference =232.2, p< 0.01), while no significant difference was observed between the Audio and Visual conditions (M difference = -0.27, p = 0.999). Additionally, the Tukey HSD post hoc test showed the SUM Model had significantly lower reaction time than the OR Model (M difference=-43.0, p< 0.05).

For Accuracy data, a similar 2 (Model: OR, SUM) × 3 (Condition: Audio, Visual, Audio-Visual) repeated-measures ANOVA showed a significant main effect of Condition, F(2, 240) = 6.80, p< 0.01, but no significant main effect of Model, F(1, 120) = 0.012, p = 0.912, and no significant Model × Condition interaction, F(2, 240) = 0.03, p = 0.97. Tukey HSD post hoc tests showed Audio-Visual conditions resulted in significantly higher accuracy than both the Audio (M difference = -0.0317, p< 0.01) and Visual conditions (M difference = -0.0322, p< 0.01), while no significant difference between unisensory conditions (M difference = -0.0006, p = 0.998).

The contour plots, Figure 5, presents the audio-visual reaction times (top row) and accuracy (bottom) for the OR Model (left) and SUM Model (right) as a function of the unisensory accumulation rates (*τ*_*A*_ and *τ*_*V*_ ). The dot-grid shows the 121 the paired unisensory accumulation rates the simulated participants. Higher values of *τ* results in faster reaction times which can impact accuracy. For both the OR Model and SUM Model faster reaction times occur for higher values of *τ*_*A*_ and *τ*_*V*_ . For the OR Model, the AV reaction times contour lines are inverted U-shaped pattern from the peak reaction time where unisensory accumulation rates are approximately the same. While for the SUM Model, the AV reaction times stay consistent along the perpendicular to the main diagonal.

**Fig 5.**
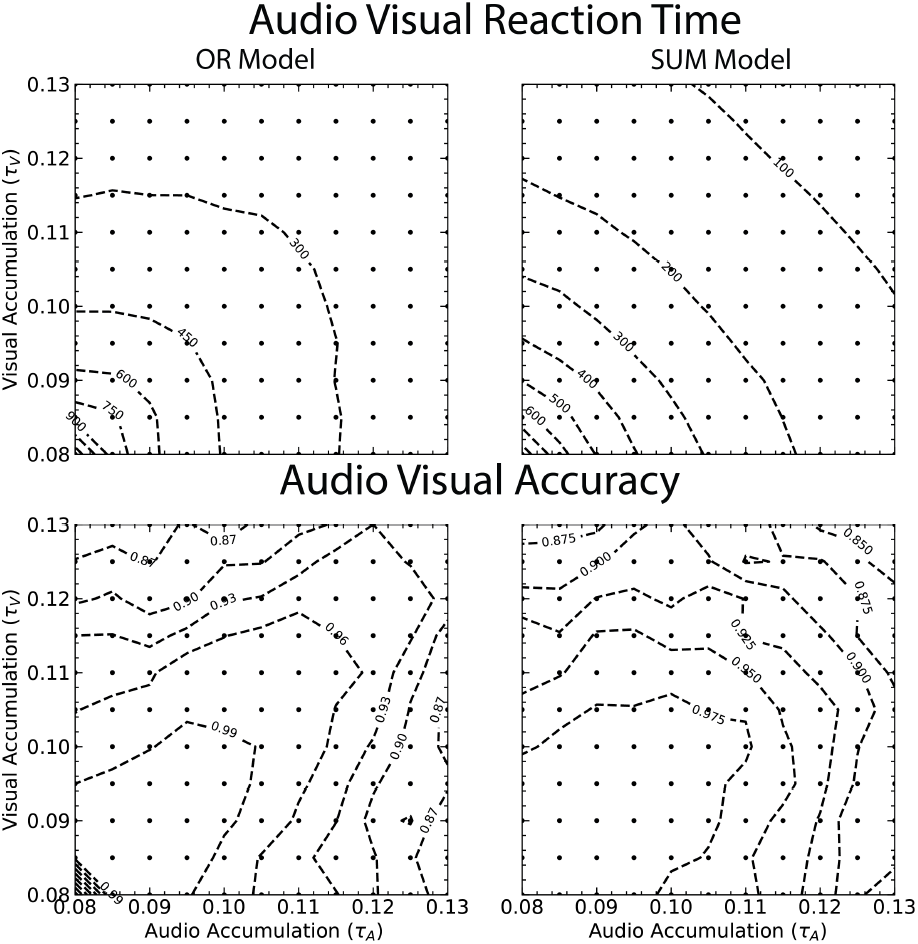
Audio-Visual Reaction Time (top) and Accuracy(bottom) Contour plots of the relationship between sensitivities to audio accumulation (*τ*_*A*_) and visual accumulation (*τ*_*V*_ ) stimuli for the OR Model (left) and SUM Model (right). The *τ* grid from Figure 1 is superimposed to identify where each of the 121 simulated participants lies. The dashed contour lines represent reaction times or accuracies. The peak of each of these contours occurs along the diagonal, where *τ*_*A*_ = *τ*_*V*_ .

Accuracy, however, declines with higher *τ* values in both models, forming an inverted U-shaped pattern along the diagonal. This suggests a speed–accuracy trade-off, where extreme *τ* values (especially when imbalanced across modalities) increase the likelihood of misses. In the lower-left corner of the OR Model accuracy plot, a sharp drop reflects trials where none of the units reached threshold within the temporal window. In these cases, responses are based on the unit with the highest partial accumulation, leading to near-chance accuracy.

#### Drift-Diffusion Analysis

A Generalized Drift-Diffusion Model (GDDM) (37) featuring free parameters of drift rate and non-decision time was fit to the reaction times and accuracy across all simulated participants. The drift rate parameter describes the rate at which evidence is accumulated, with higher drift rates correlating with faster responses. The non-decision time encompasses the duration of time excluding evidence accumulation included within the reaction time variable, incorporating motor preparation and execution processes and attentional delays arising from task or modality switching. Decreases in non-decision time correspond to faster reaction times.

For the drift rate data, a 2 (Model: OR, SUM) × 3 (Condition: Audio, Visual, Audio-Visual) repeated-measures ANOVA was conducted. The analysis revealed significant main effects of Model, F(1, 120) = 33.28, p< 0.01, and Condition, F(2, 240) = 87.20, p< 0.01, as well as a significant Model × Condition interaction, F(2, 240) = 31.94, p< 0.01. Tukey HSD post hoc tests indicated that the OR Model produced significantly higher drift rates than the SUM Model (M difference = 1.74, p< 0.01). Audio-visual conditions resulted in significantly higher drift rates than both the Audio (M difference = –4.26, p< 0.01) and Visual conditions (M difference = –4.28, p< 0.01), while no significant difference was observed between the Audio and Visual conditions (M difference = –0.02, p = 0.91).

For the non-decision time a 2 (Model: OR, SUM) × 3 (Condition: Audio, Visual, Audio-Visual) repeated-measures ANOVA was conducted. The analysis revealed no significant main effect of Model, F(1, 120) = 3.18, p=0.075, a significant effect of Condition, F(2, 240) = 61.52, p< 0.01, as well as a significant Model × Condition interaction, F(2, 240) = 3.24, p< 0.05. Tukey HSD post hoc tests showed Audio-Visual conditions resulted in significantly lower non-decision time than both the Audio (M difference = 152.1, p< 0.01) and Visual conditions (M difference =150.8, p< 0.01), while no significant difference was observed between the Audio and Visual conditions (M difference = –0.13, p = 0.996).

#### Comparison of Observed and Predicted Drift Rates

The predicted audio-visual drift rates for the multisensory audio-visual condition are calculated by fitting each participants reaction time and accuracy the drift-diffusion model to the behavioural measures of unisensory trials followed by the application of Equation 11 to calculate the predicted drift rate of an optimal observer.

Figure 6 presents the observed and predicted audio-visual drift rates for each simulated-participant the OR (cirles) and SUM (squares) models. The dashed diagonal line represents the predicted optimal audio-visual drift rate. For the OR Model, a significant positive correlation was found between observed and predicted drift rates (Spearman r(120) = 0.622, p< 0.01); however, the majority of data points fall below the diagonal, indicating sub-optimal multisensory integration. In contrast, the SUM Model showed a higher correlation coefficient (Spearman r(120) = 0.90, p< 0.01). Although most data points aligned closely with the optimal diagonal, a cluster of points with observed drift rates of 25, the max for the fitting drift parameter, fell above the predicted values, suggesting supra-optimal behavioural enhancements for a subset of simulated-participants.

**Fig 6.**
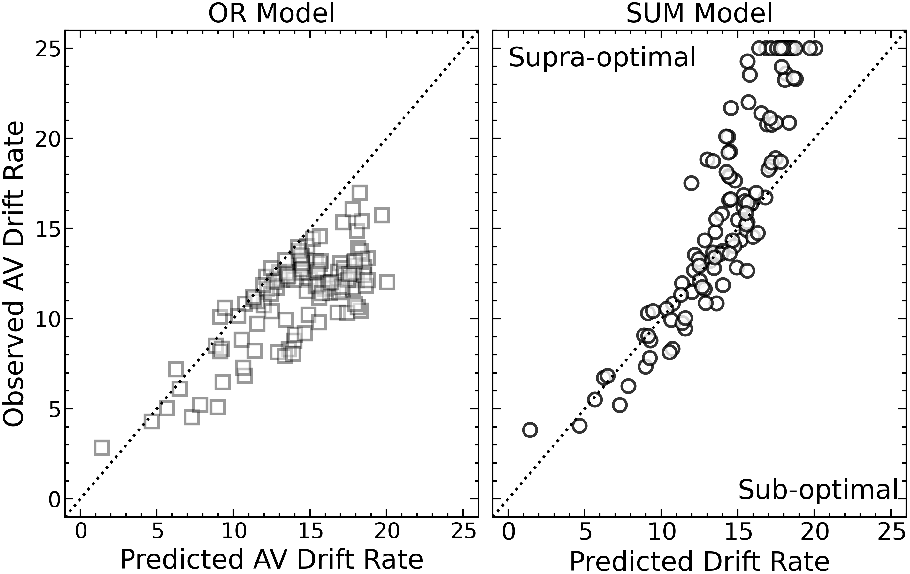
Plot for the audio-visual observed and predicted drift rate (calculated using Equation 11) for the OR Model (circles) and SUM Model (squares). The diagonal line represents optimal values of drift for the multisensory condition. Any values falling in the area below the line are sub-optimal, while any above the line are supraoptimal.

Figure 7 shows the contour colour plots of the observed audio-visual drift rates (top), predicted audio-visual drift rates (middle) and the subtraction of the observed and predicted audio-visual drift rates (bottom) as a function of the unisensory accumulation rate for the audio (*τ*_*A*_) and visual (*τ*_*V*_ ) modalities for the OR Model (left) and SUM Model (right). Notably, in the plot depicting the drift rate difference for the OR Model, darker shades of blue correspond to more negative values, indicating instances where the observed drift rate is below the predicted value, thereby suggesting sub-optimal simulated participant performance for almost all combinations of sensory accumulation rate. While for the SUM Model the difference drift rate plot shows the supra-optimal drift rates are for the higher audio and visual *τ* values.

**Fig 7.**
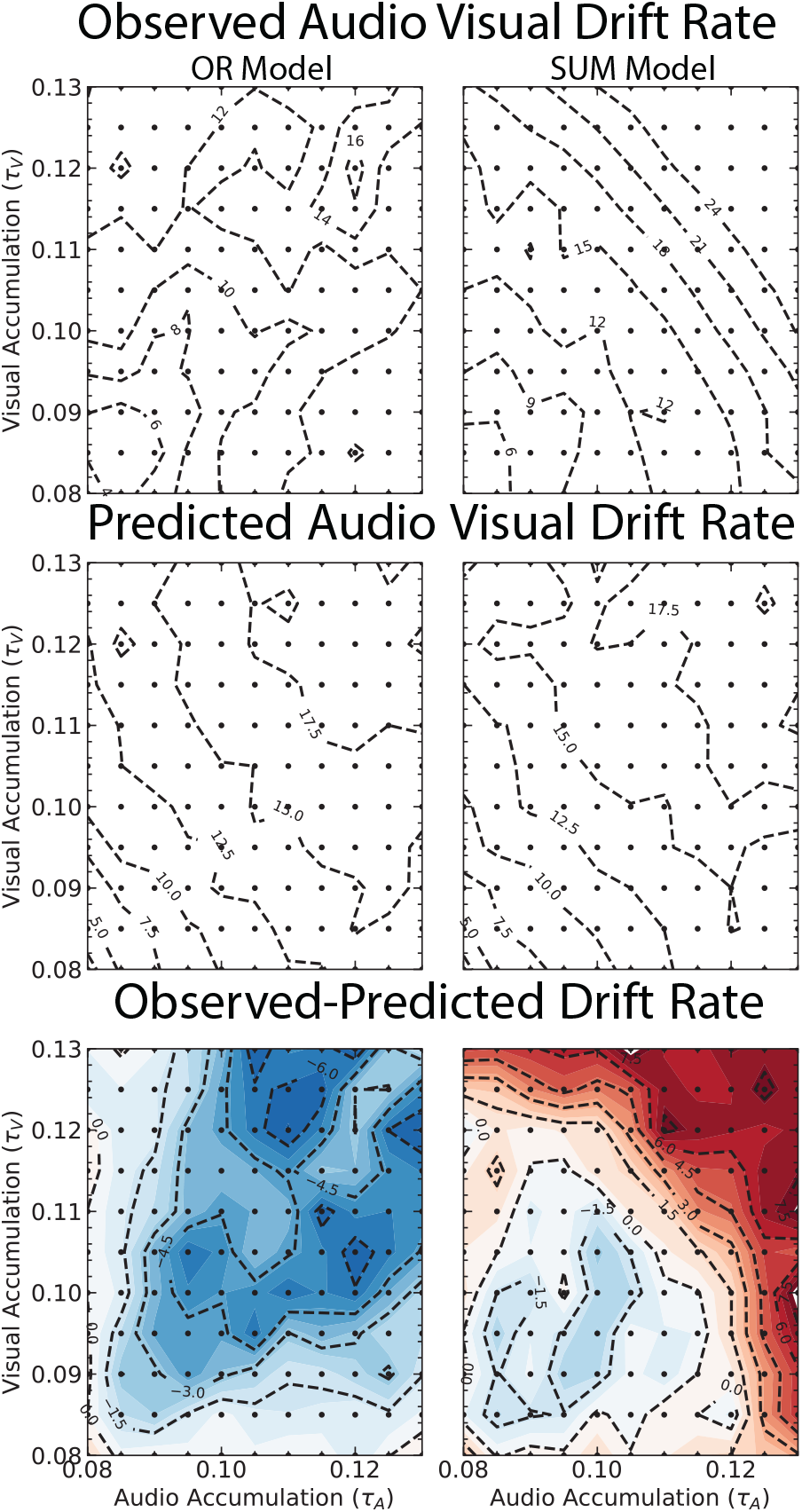
Contour plots of the relationship between audio accumulation *τ*_*A*_ and visual accumulation *τ*_*V*_ of the OR (left) and SUM (right). Models observed drift rate taken from the multisensory trials (top), the predicted drift rate calculated using the unisensory trials (middle) and the difference between the observed and predicted drift rate found by subtracting the two measures (bottom).

### Repeat Trial Winner-Take-All Model (REPEAT Model)

The widely accepted method for detecting the existence of multisensory integration in behavioural data involves identifying a deviation from winner-take-all dynamics (33). However, recent studies by (19, 23, 38) have emphasised the necessity of investigating task switching and sensory repetition effects in multisensory paradigm. When accounting for switching and repeat trial costs, the behavioural results of their study could adequately be described by winner-take-all dynamics, indicating that the behavioural results do not suggest the presence of cross-sensory facilitation or multisensory integration on this task. Their results challenge previous studies, such as (39), which reported a significant multisensory effect after adjusting reaction times to account for modality switching costs.

The adaptation of the two-variable-model by (20) presented here extends the previously discussed winner-take-all model to account for task switching (unisensory slowing) and sensory priming (multisensory speeding) effects highlighted by (19) in their audio-visual detection task. The results highlight the impact of repeating and switching costs in audio-visual trials on the behavioural outputs using a pure (blocks of A, V, or AV stimuli only) and mixed (random presentation of all three stimulus types) sensory trial design. Our winner-take-all model appears to produce some optimal measures of multisensory integration despite the model architecture specifically prohibiting this. Accounting for switch and repeat trials reveals the perceived optimality is a result of comparing speeded repeat trials to slowed switch trials, suggesting that the winner-take-all model does not account for the behavioural improvements reported experimentally. This study highlights the importance of addressing switching and priming costs associated with multisensory trials in any experimental or computational approach investigating multisensory integration. The proposed adaptation to the two-variable model aims to provide a more comprehensive understanding of the complex dynamics involved in multisensory processing.

#### Behavioural Outcomes for All Simulated Participants Across All Trials

The reaction time and accuracy data for all simulated participants across all trials are shown in Figure 8. Mean reaction time and accuracy were submitted to an analysis of variance (ANOVA), with the factors of trial type (switch, repeat) and stimulus condition: audio, visual, and audio-visual. An ANOVA showed that there were significant effects of trial type, F(1, 120) = 3.9, p< 0.05, and condition, F(2, 240) = 34.4, p< 0.01 and no significant interaction of trial type and condition F(2, 240) = 0.26, p = 0.26 for reaction time. Tukey HSD post hoc tests showed Audio-Visual conditions resulted in significantly faster reaction times than both the Audio (M difference = 175.0, p< 0.01) and Visual conditions (M difference =175.2, p< 0.01), while no significant difference was observed between the Audio and Visual conditions (M difference = 0.146, p = 0.996). Tukey HSD post hoc test showed reaction times in repeat trials were faster than switch trials (M difference = -39.44, p < 0.05). Follow-up t-tests showed that reaction times were statistically faster in the repeat trials compared with switch trials for audio (t(120)=-40.12,p < 0.01), visual (t(120)=-39.9,p < 0.01) and audio-visual conditions (t(120)=-29.2,p < 0.01).

**Fig 8.**
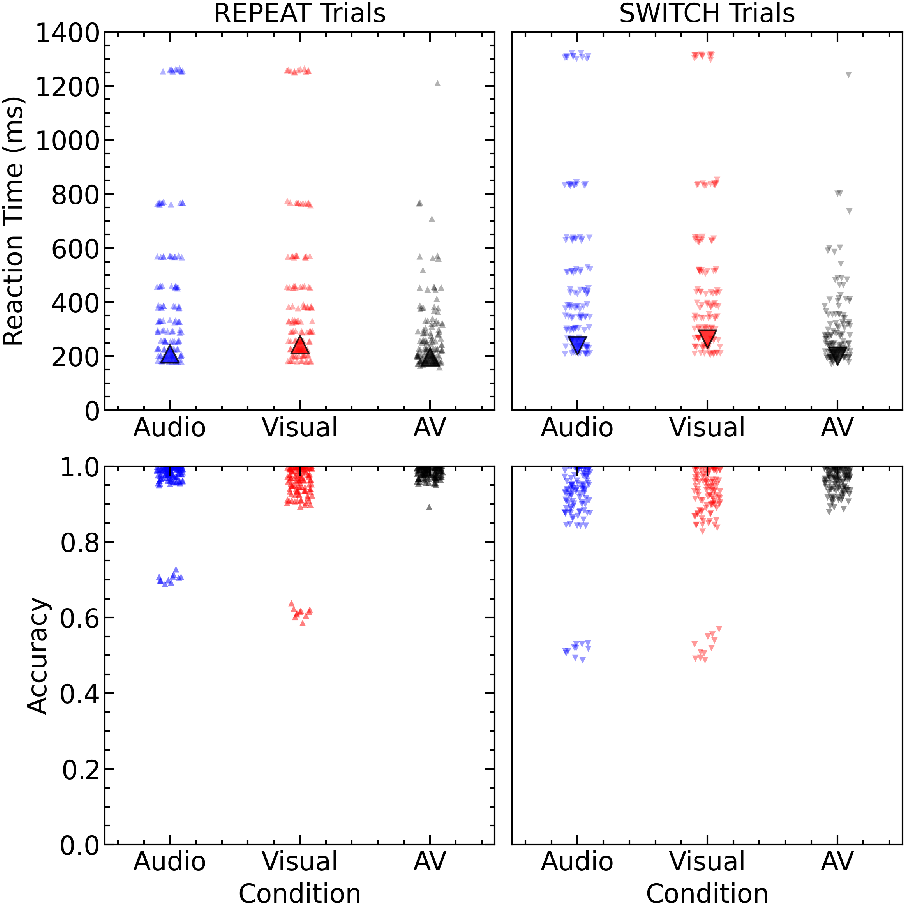
Reaction time (top) and accuracy (bottom) data for repeat trials (left) and switch trials (right). The average data for all simulated participants is shown across conditions audio (blue), visual (red) and audio-visual (grey).

Similarly, for accuracy there are significant effects of both trial types, F(1, 120) = 25.4, p< 0.01, and modality, F(2, 240) = 24.9, p< 0.01 and no significant interaction of trial type and condition F(2, 240) = 2.21, p = 0.11. Tukey HSD post hoc tests showed Audio-Visual conditions resulted in significantly higher accuracy than both the Audio (M difference = -0.043, p< 0.01) and Visual conditions (M difference =-0.058, p< 0.01), while no significant difference was observed between the Audio and Visual conditions (M difference = -0.0147, p = 0.211). Tukey HSD post hoc test showed reaction times in repeat trials were faster than switch trials (M difference = -39.44, p < 0.05). Follow-up paired t-test showed that the results for accuracy results were higher in the repeat trials compared with the switch trials for audio (t(120)=11.22,p < 0.01), visual (t(120)=11.54,p < 0.01) and audio-visual conditions (t(120)=12.57,p < 0.01).

#### Drift-Diffusion Analysis

A GDDM featuring free parameters of drift rate and non-decision time was fit for reaction times and accuracy for repeat and switch trials for each condition across all simulated participants. As illustrated in Figure 9, the drift rates are higher and the non-decision time is shorter for repeat trials than switch trials.

**Fig 9.**
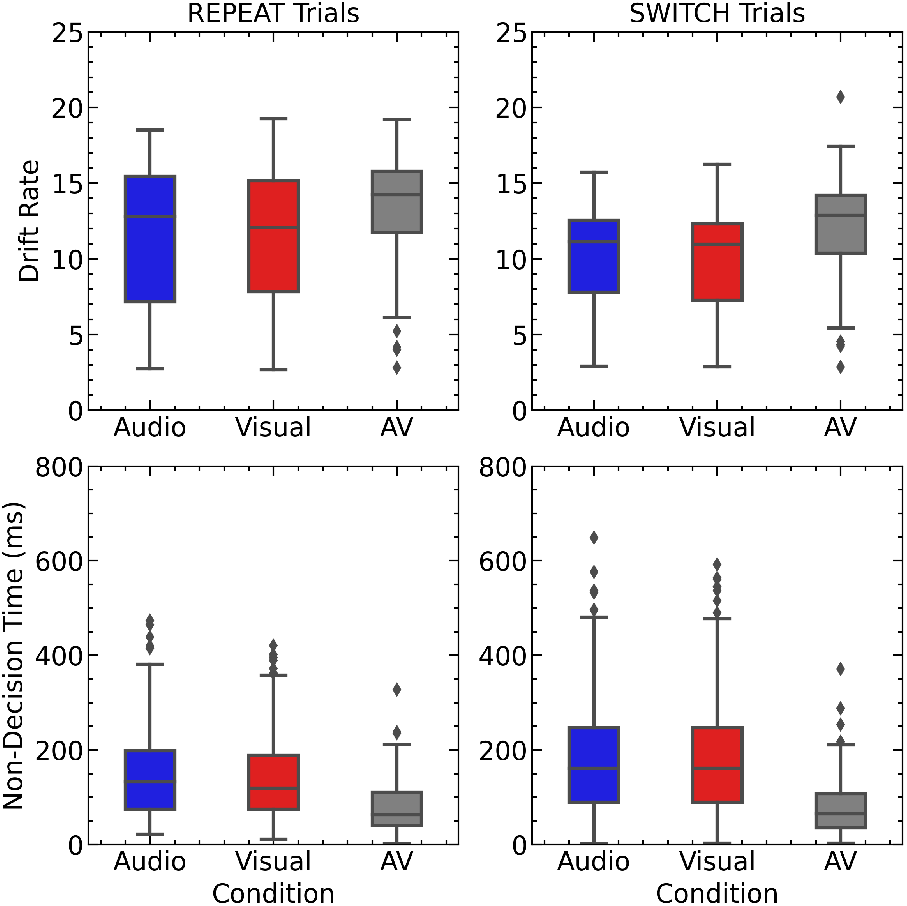
Drift rates (top) and non-decision times (bottom) as fit to data for all simulated participants on repeat trials (left) and switch trials (right). The line within each box plot is the population mean, and diamond symbols show outliers.

Drift rates and non-decision time were submitted to an analysis of variance (ANOVA), with the factors of trial type (switch, repeat) and stimulus condition: audio, visual, and audio-visual. For drift rate, an ANOVA showed that there were significant effects of trial type, F(1, 120) = 22.33, p< 0.01, and condition, F(2, 240) = 25.96, p< 0.01 and no significant interaction of trial type and condition F(2, 240) = 0.045, p = 0.95. Similarly, for non-decision time an ANOVA showed that there were significant effects of trial type, F(1, 120) = 9.73, p< 0.01, and condition, F(2, 240) = 55.65, p< 0.01 and no significant interaction of trial type and condition F(2, 240) = 2.43, p = 0.09.

#### Predicted vs Observed Drift Rates

The optimal predicted audio-visual drift rates for the multisensory repeat and switch trials are calculated using Equation 11 using the unisensory drift rates obtained after fitting the GDDM. The mean values of drift from the recorded audio, visual and audio-visual conditions, along with the predicted audio-visual drift for repeat and switch trials. On repeat trials, the observed audio-visual drift rate (M = 13.38, SD = 3.48) was significantly lower than the predicted drift rate (M = 16.69, SD = 4.4), t(120) = 7.538, p < 0.01. Similarly, on switch trials, the observed audio-visual drift rate (M = 12.15, SD = 3.1) was significantly lower than the predicted drift rate (M = 14.4, SD = 3.2), t(120) = 7.538, p < 0.01.

A scatter plot of the individual simulated participant’s data for the predicted versus observed drift rates for the audio-visual condition on repeat (“‘) and switch (‘f) trials is shown in Figure 10. The separation between repeat and switch trials is evident, with switch trials consistently tighter to or above the line of optimality.

**Fig 10.**
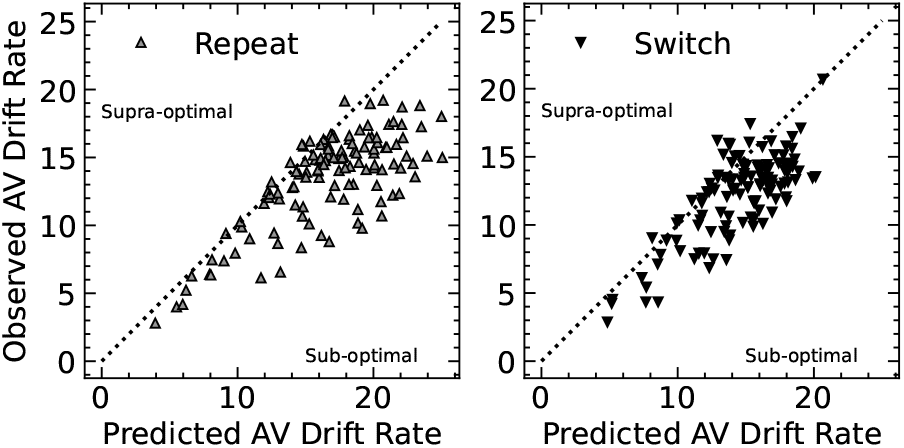
Multisensory scatter plot of predicted versus observed AV drift rates for all simulated participants on repeat (“‘) and switch (‘f) trials. The dashed line indicates optimal integration. Data points below the line indicate sub-optimal drift, while those above the line indicate supra-optimal drift.

## Discussion

We developed three models of audio-visual detection within a biologically motivated mean-field recurrent framework to test whether the multisensory speed-up in reaction times requires integration of sensory evidence. The OR Model, in which the faster of two independent unisensory processes determines the response, produced audio-visual benefits consistent with the developmental literature (6, 9) but remained sub-optimal, as expected without a combination mechanism. Where its drift rates approached the optimal prediction, this occurred in simulated participants with one markedly slower modality, where the response is essentially unisensory. The SUM Model, which linearly sums auditory and visual evidence, reached optimal integration and matched adult reaction times and accuracy at slower unisensory accumulation rates (3, 13, 22). At faster sensory accumulation rates, however, it produced supra-optimal drift rates and audio-visual reaction times faster than those observed experimentally, and near or beyond physiological limits. The REPEAT Model, an extension of the OR Model with priming on repeated modalities, reproduced the repetition effects reported by (19) and produced drift rates close to optimal on switch trials despite containing no integration. Taken together, the results suggest that neither independent processing nor full linear summation alone accounts for audio-visual detection across all participants and trial conditions. They also show that near-optimal drift rates are not sufficient evidence of integration, because trial-order effects and modality dominance can produce the same pattern.

The optimality analysis is informative only when its benchmark can discriminate between integration and independent processing, and our results show that this is not always the case. The optimal prediction is derived from the unisensory drift rates (18). When one modality is much slower than the other, the predicted optimal drift rate is therefore close to that of the dominant modality, and any model that responds on the basis of the dominant modality, such as the OR Model, will approach the optimal line without combining any evidence. This is what we observed for simulated participants with one slow unisensory accumulation rate, in both the OR and SUM Models. Proximity to optimality in this region should therefore not be read as evidence of integration. The same limitation applies to trial history. The REPEAT Model has no integration mechanism, yet its switch-trial drift rates fall closer to the optimal predictions than its repeat-trial rates, so a near-optimal drift rate can arise from priming alone. The benchmark is most diagnostic for participants with comparable auditory and visual rates, where independent processing predicts a clearly sub-optimal benefit (OR Model) and integration can reach or exceed the optimal prediction (SUM Model). Drift-diffusion analysis is therefore best used together with a race-model baseline (17) and with analyses split by trial type, rather than as a single test of integration.

The SUM Model’s main limitation is that it overshoots the optimal prediction. For participants with faster unisensory accumulation rates, its drift rates were supra-optimal and its audio-visual reaction times were faster than those observed experimentally, even though its unisensory reaction times matched the literature. Supra-optimal integration is difficult to reconcile with a linear combination of noisy cues, which sets an upper bound on the benefit, and with behavioural data, where audio-visual benefits typically fall at or below the optimal prediction (16, 18). We suggest the overshoot arises from the model’s architecture rather than from a property of integration itself. The unisensory model of (20) was designed to reproduce a single accumulation process, and its non-linear dynamics were not constrained to preserve optimality when two inputs are combined. Two modifications could test this explanation: scaling the inputs to the integrative area (for example by normalisation or weighting) (40), and imposing a lower bound on non-decision time so that the model cannot respond faster than physiological limits. If either removes the overshoot while preserving the match to adult data at slower rates, the supra-optimal responses would reflect missing constraints rather than a failure of the summation principle. If not, they would suggest that full linear summation is not a viable account of audio-visual detection at fast accumulation rates.

The REPEAT Model shows that trial-order effects alone can produce much of the audio-visual benefit that is usually attributed to integration. Priming the Hit unit of the repeated modality reproduced the faster responses on repeat trials reported by (19), and audio-visual reaction times followed their pattern, with the benefit of unisensory repeats on audio-visual trials being small. Pooling switch and repeat trials therefore blurs these effects, which can distort estimates of the multisensory benefit. This is consistent with proposals that sequence effects should be included in models of multisensory detection (16, 38). Drift rates in the REPEAT Model resembled those of the OR Model, with switch trials falling closer to the optimal prediction than repeat trials. Because the model contains no integration, this pattern is a false positive for integration, and it supports the point made above that proximity to optimality cannot by itself establish integration. The drift-diffusion fits also revealed a difference in non-decision time between repeat and switch trials for unisensory stimuli, but not for audio-visual stimuli. In the model, priming is implemented as a baseline boost to the activity of the Hit unit, which shortens the time to reach threshold rather than changing the evidence accumulation or any process outside it. That the drift-diffusion fit attributes this effect to non-decision time suggests that the parameters of a drift-diffusion model are not uniquely tied to the underlying mechanism, so priming effects can appear in non-decision time as well as in drift rate. Experimental studies using drift-diffusion analyses of audio-visual detection should therefore control for or model trial history before interpreting differences in either parameter as evidence for or against integration. Priming is also altered in schizophrenia (41) and autism (42) and changes with age, so these effects may confound group comparisons of multisensory benefit.

### Other Multisensory Models

Our framework sits at this growing intersection of neural networks that reproduce behavioural outcomes and make predictions across both domains (see (43) for review). Our framework sits between two traditions in computational studies of multisensory integration. Normative models based on Bayesian or maximum-likelihood principles predict behaviour well and give insight into computations such as causal inference (44–47), but they describe what an ideal observer would do rather than how neural circuits produce the behaviour. Circuit-level models, in contrast, specify how multisensory neurons, normalisation and connectivity could give rise to integration (40, 48–51), but they are typically constrained by single-neuron or developmental data and have been less often compared with reaction times and accuracy across populations. The mean-field recurrent model used here reproduces trial-by-trial reaction times and accuracy at the level of individual simulated participants, with parameters that can be linked to population-level neural dynamics, and its outputs can be analysed with the same drift-diffusion tools used for human data (18, 26). Within this framework, we could directly compare independent processing, summation and repetition priming, and show how they differ from an optimal observer. This is a different aim from that of the normative and circuit-level models above, and our results do not address the questions they were designed for. In particular, our models do not include causal inference or multisensory spatial or temporal binding (46), and their link to neural data is indirect: the unisensory model was constrained by recordings from LIP during visual decision-making (20, 21), not by audio-visual recordings. Machine learning approaches to integration (52–54) can capture complex input statistics but are harder to link to specific circuit mechanisms. A natural next step is to combine our framework with these approaches, for example by adding a causal inference stage that decides whether auditory and visual signals should be integrated or kept separate.

#### Limitations

Several limitations should be considered when interpreting these results. First, the models were compared with published group-level data rather than fitted to individual participants, so the match to behaviour is qualitative and cannot establish which model best describes any one person. Second, the range of unisensory accumulation rates and the number of simulated participants (121) were chosen to span plausible human behaviour rather than estimated from data, so the proportion of participants showing supra-optimal or near-optimal behaviour depends on this choice and should not be read as a population estimate. Third, we simulated speeded detection of congruent, salient stimuli only. The models therefore make no predictions about conflicting or weak stimuli, spatial or temporal binding, or causal inference, and our conclusions may not extend to decision-making tasks. Fourth, the unisensory model was developed for visual decision-making in LIP (20), and its adaptation to audio-visual detection is an assumption, not something constrained by audio-visual recordings. Fifth, the OR, SUM and REPEAT Models represent discrete hypotheses. Real behaviour may lie between them, or combine integration with repetition effects, which we did not model. Finally, the optimality analysis depends on the drift-diffusion fits, and as discussed above, parameters such as drift rate and non-decision time may not map uniquely onto model mechanisms.

#### Conclusion

We simulated three models of audio-visual detection in a biologically motivated recurrent framework to ask whether the multisensory speed-up requires integration of sensory evidence. Independent processing (OR Model) produced improved reaction times and accuracy consistent with developmental data but sub-optimal drift rates. Linear summation (SUM Model) reached optimal integration at slower unisensory accumulation rates but produced supra-optimal, biologically implausible responses at faster rates. Modality repetition (REPEAT Model) reproduced priming effects and near-optimal drift rates on switch trials without any integration. Together, these results show that near-optimal drift rates are not sufficient evidence of integration, and that trial history and modality dominance can mimic it. They also suggest that neither independent processing nor full linear summation accounts for audio-visual detection across all participants and conditions. These findings yield testable predictions. If repetition priming contributes to the apparent multisensory benefit, the benefit measured in audio-visual detection should shrink when trial history is controlled or modelled, and group differences in priming should alter group differences in measured integration. Future models should test whether a mechanism that limits summation, such as normalisation or a physiological lower bound on non-decision time, removes the supra-optimal responses while preserving the match to adult data, and whether a model that combines integration with repetition priming accounts for behaviour better than any single model. Extending the framework from detection to decision-making, and adding a causal inference stage, would test whether the conclusions hold when stimuli conflict or signals originate from different sources. By linking mechanisms to behaviour in one framework (43), this approach provides a way to separate the contributions of integration, independence and repetition to multisensory benefits across development, ageing and clinical populations.

#### Conclusions

The models successfully replicated established behavioural measures of audio-visual detection across various human populations. The OR Model generated sub-optimal behavioural enhancements, consistent with those observed in young children (6). The SUM Model yielded behavioural improvements that aligned with those seen in healthy adults with lower cue sensitivity, as well as in individuals with Parkinson’s disease and age-matched healthy controls (22). Analysis using the REPEAT model revealed that ignoring the behavioural benefits of modality repetition can obscure the advantages of multisensory trials over unisensory ones (19). Each model’s ability to reproduce distinct behavioural outcomes demonstrates the utility of this approach in testing and replicating potential multisensory mechanisms documented in the literature as well as testing different metrics for behavioural multisensory integration like the drift diffusion approach.

## Acknowledgements

The authors used Claude (Anthropic, Sonnet 5.5) to assist with editing and rewriting portions of this manuscript for clarity and language refinement. All intellectual content and final edits were made by the authors. The research carried out in this publication was funded by the Irish Research Council under grant number GOIPG/2020/943 awarded to R. M. Brady.

## Citation Diversity Statement

Recent work in several fields of science has identified a bias in citation practices such that papers from women and other minority scholars are under-cited relative to the number of such papers in the field (55–63). First, we obtained the predicted gender of the first and last author of each reference by using databases that store the probability of a first name being carried by a woman (59, 64). By this measure (and excluding self-citations to the first and last authors of our current paper), our references contain 9.83% woman(first)/woman(last), 19.92% man/woman, 19.63% woman/man, and 50.62% man/man. This method is limited in that a) names, pronouns, and social media profiles used to construct the databases may not, in every case, be indicative of gender identity and b) it cannot account for intersex, non-binary, or transgender people. Second, we obtained predicted racial/ethnic category of the first and last author of each reference by databases that store the probability of a first and last name being carried by an author of colour (65, 66). By this measure (and excluding self-citations), our references contain 12.24% author of colour (first)/author of colour(last), 14.35% white author/author of colour, 16.54% author of color/white author, and 56.86% white author/white author. This method is limited in that a) names and Florida Voter Data to make the predictions may not be indicative of racial/ethnic identity, and b) it cannot account for Indigenous and mixed-race authors, or those who may face differential biases due to the ambiguous racialization or ethnicisation of their names.

